# Plant PAXX has an XLF-like function and stimulates DNA end-joining by the Ku-DNA ligase IV-XRCC4 complex

**DOI:** 10.1101/2023.05.26.542399

**Authors:** Hira Khan, Takashi Ochi

**Affiliations:** The Astbury Centre for Structural Molecular Biology, School of Molecular and Cellular Biology, Faculty of Biological Sciences, University of Leeds, Leeds LS2 9JT

## Abstract

Non-homologous end joining (NHEJ) plays a major role in repairing DNA double-strand breaks (DSBs) and is key to genome stability and editing. The minimal core NHEJ proteins, namely Ku70, Ku80, DNA ligase IV and XRCC4, are conserved, but other factors vary in different eukaryotes groups. In plants, known NHEJ proteins are the core factors only, and the molecular mechanism of plant NHEJ remains unclear. Here, we report a previously unidentified plant ortholog of PAXX, the crystal structure of which showed a similar fold to human PAXX. However, plant PAXX has similar molecular functions to human XLF, by directly interacting with Ku70/80 and XRCC4. This suggests that plant PAXX combines the roles of mammalian PAXX and XLF and that these functions merged into a single protein during evolution. This is consistent with a redundant function of PAXX and XLF in mammals.

## Introduction

Non-homologous end joining (NHEJ) is a major pathway to repair DNA double-strand breaks (DSBs)^1^. NHEJ is evolutionarily conserved between bacteria and eukaryotes, and is comprised of two core proteins, Ku and DNA ligase^2^. In eukaryotes, most organisms have minimal core proteins; Ku70/80 heterodimer and DNA ligase IV/XRCC4 (LIG4/XRCC4) heterotrimer. Other NHEJ-specific proteins, particularly in mammals, include DNA-dependent protein kinase catalytic subunit (DNA-PKcs), XRCC4-like factor (XLF), Paralog of XRCC4 and XLF (PAXX), Artemis, X-family DNA polymerases (POL), Aprataxin-like protein (APLF) and cell cycle regulator of non-homologous end joining (CYREN) / the modulator of retrovirus infection (MRI)^1^. During evolution, NHEJ gained complexity by incorporating more proteins to maintain genome stability in order to cope with challenges during the development of vertebrates. NHEJ is also important for genome editing, for instance, by CRISPR-Cas9 because it introduces DSBs by enzymes. The current understanding of eukaryotic NHEJ mostly comes from studies in yeasts and mammals. However, our knowledge of NHEJ may not be directly applied to different organisms because they often have different sets of NHEJ proteins^3^.

Plants provide examples of such organisms, which have Ku70, Ku80, LIG4 and XRCC4^4–6^ but lack apparent orthologs of other core proteins such as DNA-PKcs and XLF. Plant NHEJ proteins play important roles in DSB repair, telomere maintenance, genome editing *via* CRISPR-Cas9, as observed in other organisms, and also in the stable integration of *Agrobacterium* transfer DNA (T-DNA) into plant genomes^7–9^. A number of studies in mammalian NHEJ highlighted the importance of DNA-PKcs and XLF in DNA-end synapsis^10– 21^. Although yeasts do not have DNA-PKcs, they have XLF orthologs Nej1^22,23^, which play a similar role to XLF. This raises a question of how plant NHEJ works only with Ku70/80 and LIG4/XRCC4 complexes at the molecular level.

Here, our evolutionary analysis of NHEJ proteins identified the plant ortholog of PAXX. The crystal structure of *Arabidopsis thaliana* PAXX (AtPAXX) shows that it has a similar fold to human PAXX and interacts with Ku70/80 and, unlikely human PAXX, with XRCC4. *In vitro*, these interactions stimulated DNA end-joining by the LIG4/XRCC4 complex. Thus, our results suggest that AtPAXX plays a hybrid role in human XLF and PAXX.

## Results

### Overview of the conservation pattern of NHEJ proteins

We searched for amino-acid sequences of known NHEJ proteins to understand their conservation in eukaryotes. A broad group of eukaryotes have Ku70, Ku80, DNA-PKcs, DNA ligase IV, XRCC4, XLF, PAXX, Artemis, POL including polμ/λ/TdT, APLF and CYREN/MRI (Figure 1 and Table S1). CYREN/MRI is the least conserved protein. Sea urchin has polynucleotide kinase-phosphatase-like protein fused to the motif that is similar to the N-terminal Ku-binding motif of CYREN/MRI (Table S1). APLF orthologs are also limited to multicellular organisms but are present more broadly than those of CYREN/MRI. A choanoflagellate *Salpingoeca rosetta* has an APLF-like protein possessing a poly (ADP-ribose)-binding zinc finger (PBZ) domain, but the protein also has a BRCT domain and core domains of DNA ligase (Table S1). In most opisthokonta and ciliates, the full set of NHEJ proteins except for Artemis, APLF and CYREN/MRI are present. *C. elegans* has a unique DSB-repair system among multicellular organisms because it has Ku70, Ku80 and LIG4 orthologs but lacks XRCC4 orthologs (Figure 1). A structural homolog search using the DALI server against AlphaFold models^24^ of *C. elegans* proteins yields two DNA ligases; DNA ligase I and IV (accession numbers: Q27474 and Q95YE6 respectively). The latter has one BRCT domain like DNA ligase III^25^ and lacks the XRCC4-interacting motif^26^ in line with the lack of XRCC4. Apart from Ku70/80 and DNA-PKcs, core NHEJ proteins are absent in the majority of the organisms belonging to the excavata group. A recent evolutional analysis of DNA-PKcs identified its orthologs in a wide range of eukaryotes including *Naegleria*^27^ and *Leishmania major*^28^. In addition, we found that *Trypanosome cruzi* has a putative DNA-PKcs ortholog that does not have the FAT, kinase and FATC domains but the heat-repeat domain, of which fold is predicted to be similar to that of human DNA-PKcs (Figure 1 and Figure S1B). Note that we could not identify a similar DNA-PKcs ortholog in *Trypanosome brucei*. Uniquely, as reported previously^29^, core NHEJ proteins are absent in *Giardia lamblia* (Figure 1).

**Figure 1.**
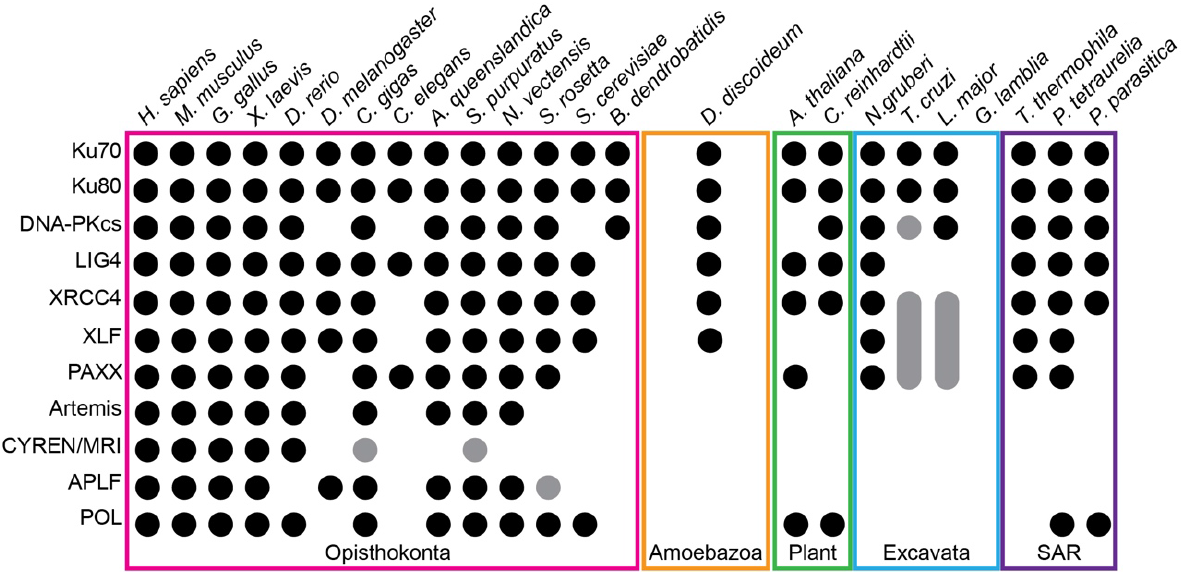
Evolutionarily conserved NHEJ proteins in eukaryotes. A diagram summarises the presence of NHEJ proteins in various groups of eukaryotes. Black circles represent the presence of NHEJ proteins with confidence, whereas grey circles represent uncertainty.

Some of the end-processing enzymes in NHEJ also work in different DNA-repair pathways apart from Artemis and DNA polymerases (μ, λ and TdT). Therefore, we here discuss the evolutional analysis of these proteins only. Metallo-β-Lactamase/β-CASP domains containing proteins can be found in all kingdoms and work in DNA and RNA processing^30^. Among them, a unique structural feature of Artemis is a zinc-finger motif (H228, H254, C256 and C272) in the β-CASP domain^31^. Metallo-β-Lactamase/β-CASP domains-containing proteins having the zinc-finger motif were limited to jawed organisms (Figure 1). X-family DNA polymerases that are specific to NHEJ carry BRCT domains^32^. Outside opisthokonta, such polymerases are found in plants^33^ and ciliates (Figure 1).

Altogether, our evolutional analysis suggested that Ku70/80, DNA-PKcs, LIG4, XRCC4, XLF and POL are core NHEJ proteins present in both unicellular and multicellular eukaryotes.

### PAXX is conserved among uni- and multi-cellular organisms

We had previously suggested that PAXX orthologs were present in Filozoa excluding insects and nematodes^34^. Further sequence mining identified the orthologs in *C. elegans*, amoeboflagellates, ciliates, excavates and plants (Figure 1). We found that the recently identified NHEJ factor in *C. elegans* NHJ-1^35^ is likely to be a PAXX ortholog because of its similarity to the PAXX fold (Figure S1A). In amoebazoa, PAXX is not present in *Dictyostelium discoideum* but is present in other amoebozoa such as *Acanthamoeba castellanii* and *Planoprotostelium fungivorum* (accession numbers: L8H683 and A0A2P6NT14). Ciliates such as *Tetrahymena* and *Paramecium* have PAXX orthologs (Figure 1). Based on sequences of the ciliate PAXX, we found that the orthologs had been known as Die5p (Defective in internal eliminated sequence (IES) excision 5)^36^. Ciliates have complex genomic structures and have two nuclei, called the macronucleus and micronucleus. During macronucleus differentiation, DNA rearrangements occur by eliminating IESs from chromosomes^37^, and generated DSBs are joined by NHEJ. Die5p was proposed as a DSB-repair protein working downstream of Ku80^38^, which is consistent with the role of PAXX in humans. Interestingly, *Trypanosome cruzi* has two putative XRCC4-like proteins that are neither centrosomal protein SAS6^39,40^ nor CCDC61^41^ and have phosphodiesterases at the C-termini (Table S1 and Figure S1C). A similar domain arrangement was also found in *Leishmania*, implying this phosphodiesterase fusion is conserved in those organisms. *Naegleria gruberi*, another organism belonging to excavata, has most of the core NHEJ proteins including XLF and PAXX (Figure 1).

These results suggest that PAXX was an NHEJ factor in the early eukaryotes but probably limits its function to a certain type of DSBs at least in mammals due to redundancy with other proteins such as XLF^42–47^.

### Plant NHEJ lacks XLF but has PAXX

The plant is the only group that does not have XLF but PAXX in the organisms with LIG4/XRCC4 shown in Figure 1. This implies that PAXX might play a major role in plant NHEJ. We therefore investigated the structure and function of plant PAXX.

Plant PAXX is well-conserved among streptophyta (Table S2) and has a putative Ku-binding site at the C-terminus, which is similar to PAXX orthologs (Figure 2A). Since the XLF C-terminus also interacts with Ku^48^, plant PAXX is still possibly the ortholog of XLF. An amino acid sequence analysis of PAXX and XLF orthologs indicated that AtPAXX is more similar to the PAXX family than the XLF family (Figure 2B). To define whether plant PAXX is the human PAXX ortholog, we determined the crystal structure of full-length *A. thaliana* PAXX (AtPAXX) at 3.6-Å resolution (Table 1). AtPAXX has a short coiled-coil domain that mediates homodimerisation as observed in the structure of human PAXX (Figure 2C) and is more similar to human PAXX than XLF (Figure 2D): Unlike XLF, the alpha-helical region does not fold back to the coiled-coil domain, confirming that this AtPAXX is not an ortholog of XLF but that of PAXX. As observed in the crystal structure of full-length human PAXX^34^, we were unable to see the electron density of residues after ∼150, indicating that they were disordered *in crystallo*. The lack of XLF orthologs in *A. thaliana* was further confirmed by searching *A. thaliana* proteins that have a similar fold to XRCC4 using DALI server against AlphaFold models^24^, which identified *A. thaliana* XRCC4 (AtXRCC4) and AtPAXX only. These results suggest that plants have orthologs of PAXX but not those of XLF.

**Table 1.**
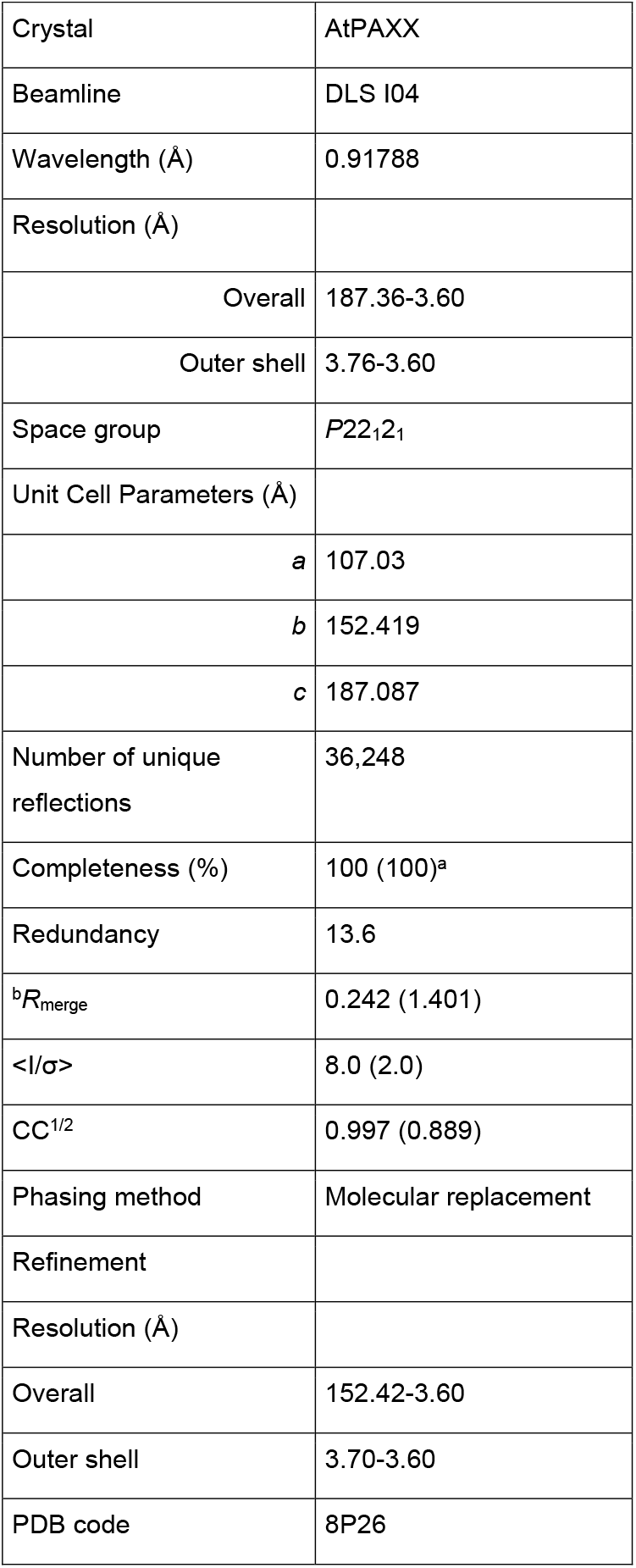
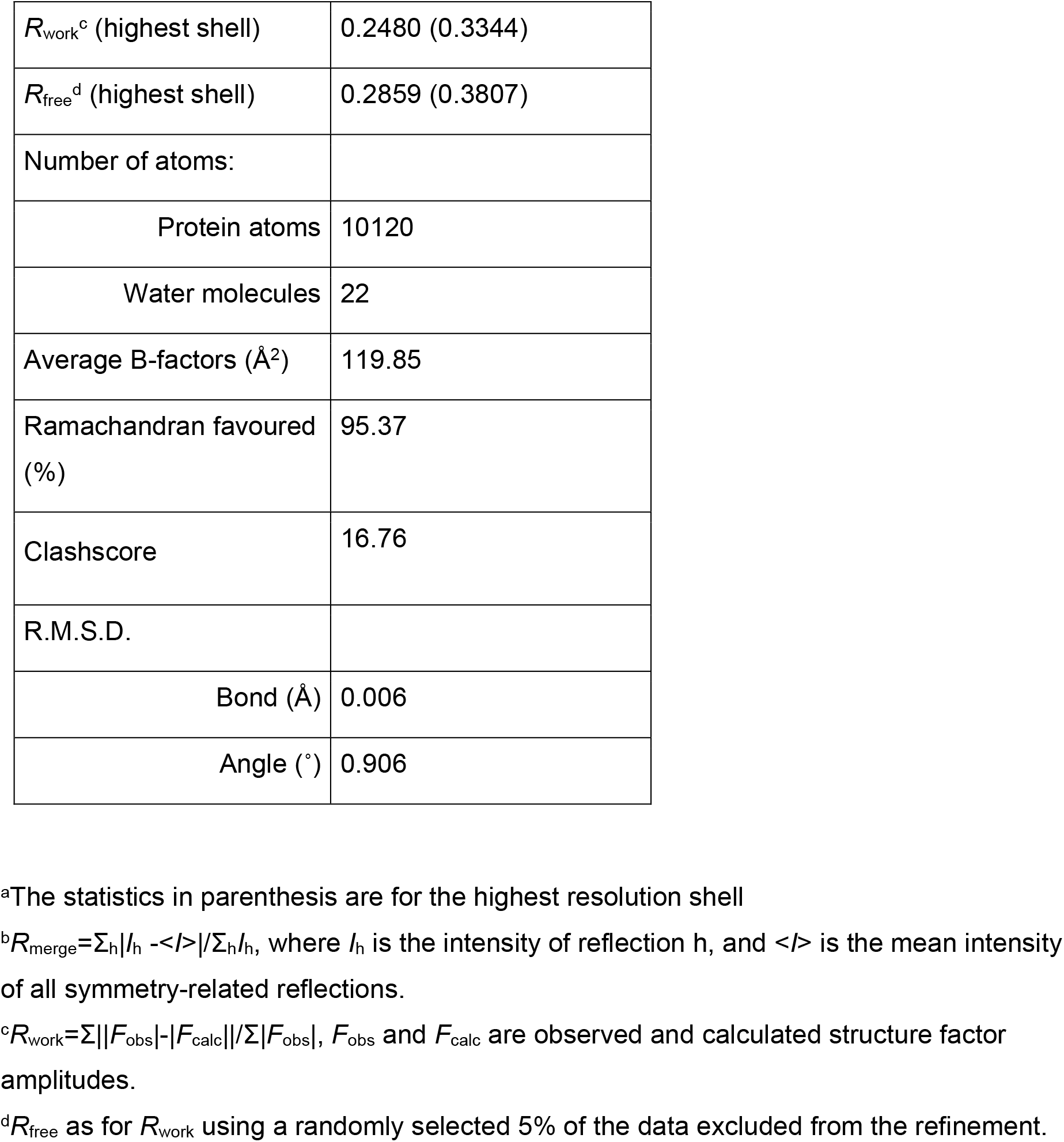
Data collection, phasing and refinement statistics of the CCDC61 crystal structures.

**Figure 2.**
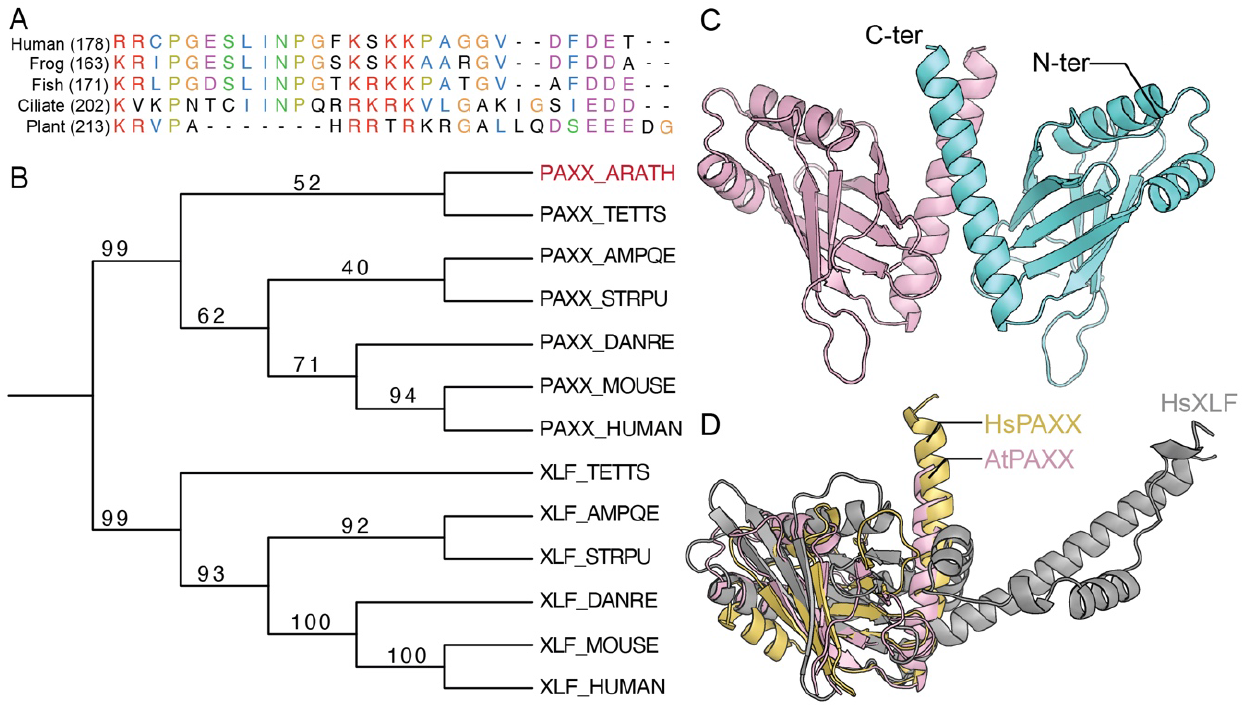
Plants have orthologs of PAXX. (A) Sequence alignment of the C-termini of PAXX orthologs. Numbers in brackets are residue numbers. (B) Phylogenetic tree of PAXX and XLF orthologs. The phylogenetic tree was generated by aligning sequences of PAXX and XLF orthologs from human (HUMAN), mouse (MOUSE), zebrafish (DANRE), sea urchin (STRPU), sponge (AMPQE), *Tetrahymena* (TETTS) and AtPAXX (ARATH in red). Numbers on the three are bootstrap values (*n*=100). (C) Crystal structure of AtPAXX dimer. Two chains are shown in a cartoon representation with two different colours. The electron density of residues ∼150-236 was not observed due to the flexibility of the C-terminal region of AtPAXX. (D) Structural comparison of AtPAXX with human PAXX and XLF. The head domain of the structure of AtPAXX was superimposed on those of human PAXX and XLF (PDB codes: 3WTD and 2QM4).

### Plant PAXX interacts with Ku and XRCC4

Human PAXX interacts with Ku, which is essential for the PAXX function^34^. Since AtPAXX has a similar motif to the Ku-interacting motif of human PAXX (Figure 2A), we tested whether AtPAXX interacts with *A. thaliana* Ku70/80 (AtKu) by a GST-pulldown experiment using recombinant AtKu expressed in insect cells^49^. The result showed that AtPAXX, particularly its C-terminus (residues 190-236), interacts with AtKu (Figure 3A). The interaction was further confirmed by an electrophoretic mobility shift assay (EMSA), which showed that bands of the AtKu-DNA complex were further shifted by adding full-length AtPAXX but not its truncated construct (residues 1-154) (Figure 3B). An AlphaFold model of the AtPAXX-Ku complex indicated that L228 of AtPAXX plays a key role in the interaction (Figure 3C). We therefore mutated the leucine to glutamate and showed that the L228E mutation abolished the interaction (Figure 3B). These suggested that AtKu interacts with AtPAXX via its C-terminal region, particularly L228.

**Figure 3.**
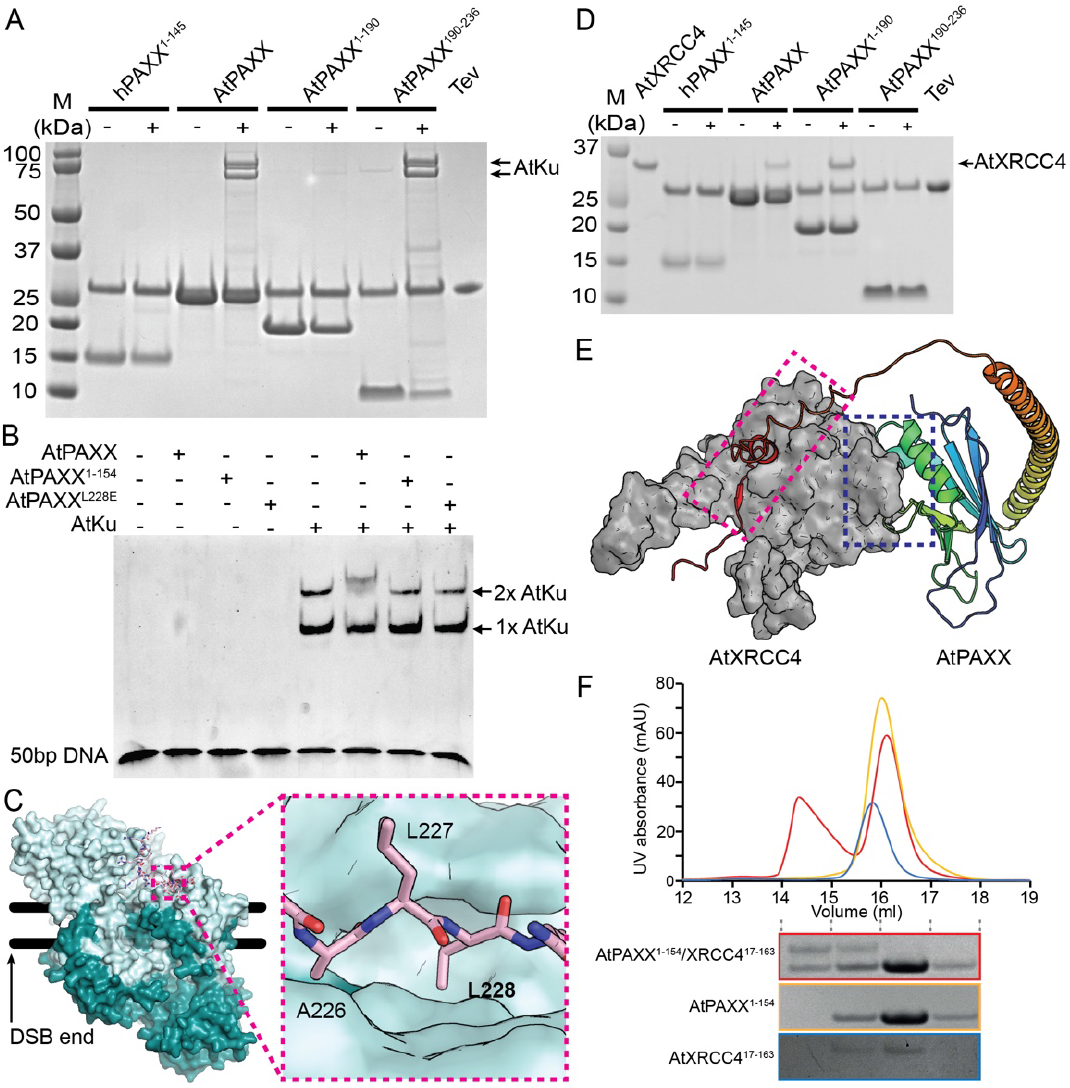
AtPAXX interacts with AtKu and AtXRCC4. (A) The C-terminus of AtPAXX interacts with AtKu. An SDS-PAGE gel shows the result of a GST-pulldown experiment. “+” indicates GST-tagged PAXX constructs were incubated with insect cell lysates expressing AtKu. The PAXX constructs were released from glutathione beads by digesting the linker between GST and PAXX by Tev. (B) AtPAXX interacts with AtKu bound on DNA. An EMSA result shows that AtPAXX but not AtPAXX^1-154^ and AtPAXX^L228E^, interacts with mainly two AtKu molecules (2x AtKu) bound on 50bp DNA. (C) AlphaFold model of AtPAXX/Ku complex. The model shows that the C-terminus of AtPAXX interacts (pink) with the surface predominantly created by AtKu70 (cyan; AtKu80 is shown in deep teal). Two black lines represent double-stranded. (D) AtPAXX interacts with AtXRCC4. An SDS-PAGE gel shows the result of a GST-pulldown experiment. “+” indicates GST-tagged PAXX constructs were incubated with AtXRCC4. The experiment was done in a similar manner to (B). (E) AlphaFold model of AtPAXX/XRCC4 complex. Potential-interacting regions of AtPAXX and AtXRCC4 (grey) were indicated with blue (N-terminal interaction) and red (C-terminal interaction) dotted rectangles. AtPAXX is presented in a gradient of blue (N-terminus) and red (C-terminus). (F) The N-terminal domain of AtPAXX interacts with that of AtXRCC4. A graph shows size-exclusion chromatography analyses of the complex and individual proteins (red) of AtPAXX (yellow) and AtXRCC4 (blue). SDS-PAGE gels show four fractions between elution volumes between 14 and 18 ml of the complex and individual samples.

We speculated that AtPAXX might interact with AtXRCC4 in a similar manner to the interaction of human XRCC4 and XLF. We therefore purified AtXRCC4 and tested for its interaction with AtPAXX by a GST-pulldown assay. The result showed that AtXRCC4 interacts with AtPAXX (Figure 3D). An AlphaFold model of the AtPAXX/XRCC4 complex predicted that both N- and C-termini of AtPAXX could interact with AtXRCC4 (Figure 3E). However, AtXRCC4 did not interact with the C-terminal low complexity region (residues 190-236) but with the folded region (residues 1-190) of AtPAXX. We further confirmed this result by the size exclusion chromatography using truncated constructs of AtPAXX (residues 1-154; AtPAXX^1-154^) and AtXRCC4 (residues 17-163) (Figure 3F). Thus, these results support our model of AtPAXX interacting with AtXRCC4 *via* their head domains.

Altogether, we have biochemically shown that AtPAXX interacts with AtKu and AtXRCC4 and that AtPAXX has functions equivalent to the human XLF protein.

### Plant PAXX stimulates end joining by DNA ligase IV

We next tested whether AtPAXX could stimulate the end joining by AtLIG4/XRCC4 *in vitro*. Blunt-ended linearised plasmids were incubated with different combinations of AtNHEJ proteins. The results showed that the end joining of AtLIG4/XRCC4 was inefficient but was slightly stimulated by AtKu and further by adding AtPAXX (Figure 4A). AtPAXX^L228E^, which does not interact with AtKu (Figure 3B), also stimulated the end-joining by AtLIG4/XRCC4 in the presence of AtKu but ∼1.4-fold less efficient than the wild type AtPAXX (Figure 4A), indicating that the interaction between AtPAXX and AtXRCC4 is likely to play a key role in the end-joining but that between AtPAXX and AtKu is also important. However, the end joining barely occurred without AtKu, which plays a central role in bridging two DNA ends.

**Figure 4.**
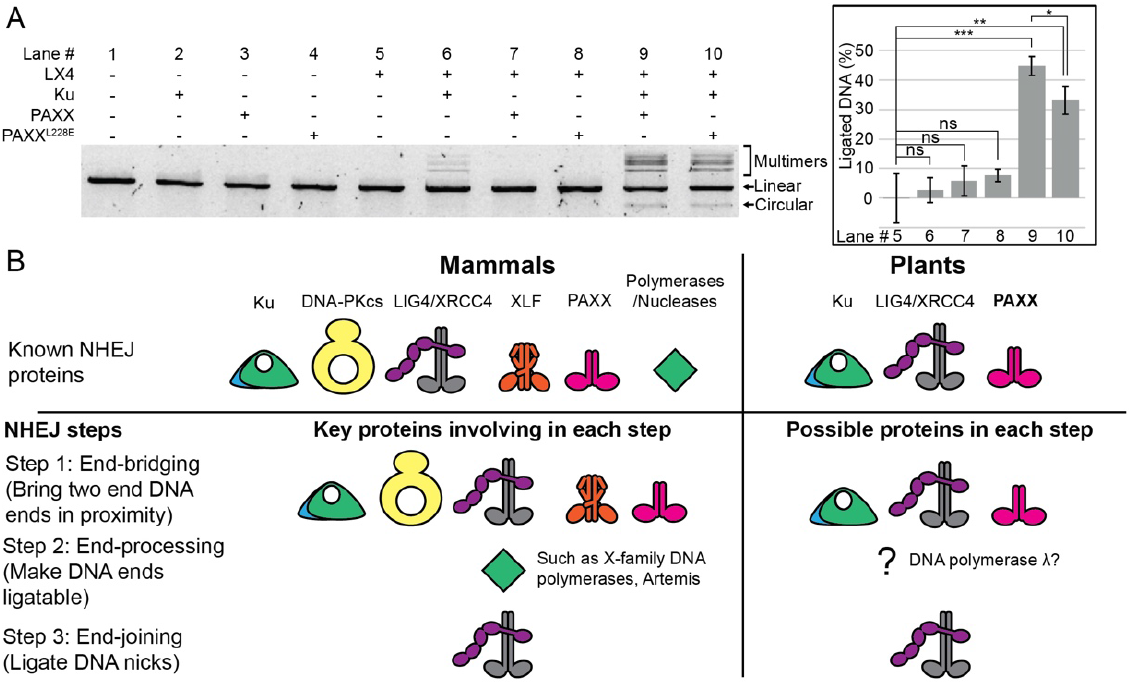
AtPAXX is a plant NHEJ factor. (A) AtPAXX stimulates blunt-ended DNA-joining by AtKu and AtLIG4/XRCC4. An agarose gel shows the result of a DNA ligation assay. Linearised blunt-ended plasmids (linear) were incubated with different protein combinations indicated by “+”. An arrow and bracket indicate ligated DNA substrates (circular and multimers). The bar graphs on the right show gel densitometry analysis results (*n*=3). Bars and error bars are averages and standard deviations. *p*-values of *t*-test results calculated against the result of AtLIG4/XRCC4 only (lane 4) are indicated by ns, *, ** and *** (*p* >0.05, 0.05, ≤0.01 and ≤0.001 respective). (B) Comparison of mammalian and plant NHEJ. Protein icons indicate known NHEJ proteins involved in three different steps.

Collectively, these results show that AtPAXX interacts with both AtXRCC4 and AtKu, and suggest that these interactions are important for DNA end-joining by AtLIG4/XRCC4.

## Discussion

We showed evolutional conservation of core NHEJ proteins, Ku70, Ku80, DNA-PKcs, LIG4, XRCC4, XLF, PAXX and POL and that plants have PAXX orthologs that interact with both Ku70/80 and XRCC4, and stimulate DNA end-joining by LIG4/XRCC4. Thus, plant PAXX orthologs play a hybrid role equivalent to human PAXX and XLF in NHEJ (Figure 4B). Thus, through our evolutional analysis of NHEJ proteins, we discovered previously unidentified PAXX orthologs in plants.

NHEJ is conserved between prokaryotes and eukaryotes. Bacterial NHEJ utilises Ku homodimer and DNA ligase D, which is a multi-functional single-polypeptide comprised of DNA ligase, polymerase and phosphoesterase domains^2^. Its exact molecular mechanism remains to be elucidated, but these proteins are likely to carry out three NHEJ steps similar to those in mammals. The minimum core NHEJ complexes in eukaryotes are Ku70/80 and LIG4/XRCC4. LIG4 is essential for NHEJ and works only in the pathway. However, kinetoplastids lack the ligase while having Ku70/80 and putative DNA-PKcs orthologs and XRCC4-like proteins (Figure 1). In those organisms, Ku is not important for NHEJ but for telomere maintenance^50^, a known conserved function of DNA-PK^51^. Interestingly, the putative DNA-PKcs orthologs of *Trypanosome cruzi* possess the HEAT-repeat domain only (Figure S1) and therefore might play roles in DNA-end bridging and protection, which are non-kinase functions of DNA-PKcs^18,20,21,52^. Since these functions are regulated by the kinase activity of DNA-PKcs in mammals^53^, other kinases might control DNA-PK in *Trypanosome cruzi*. Thus, this analysis highlights that NHEJ and its constituent proteins seem to have evolved differently to adapt to different eukaryotes.

A unique feature of eukaryotic NHEJ is the XRCC4 superfamily, including XRCC4, XLF and PAXX. These scaffold proteins play crucial roles in NHEJ by mediating protein-protein interactions. Since XRCC4 has an essential role in the stability of LIG4^54,55^, these proteins are almost always present as a pair in eukaryotes except for *C. elegans* (Figure 1). XLF and PAXX have a redundant function^42–47^, and PAXX is often absent in eukaryotic organisms (Figure 1). We showed that plants do not possess XLF but PAXX orthologs. How can we conclude that the orthologs are not XLF but PAXX? We concluded this because their structures are more similar to human PAXX than XLF. One distinguishing feature in the human proteins is the relative angle of the head and coiled-coil domain, with XLF different from XRCC4 and PAXX^34^. The angle of AtPAXX is similar to human PAXX and unlike that of XLF. Also, XLF has a fold-back α-helical structure, which was not observed in AtPAXX. Based on the observation that AtPAXX interacts with both XRCC4 and Ku so does XLF, we consider that PAXX and XLF might diverge from a common origin during evolution.

Recent cryo-EM structures of the short-range NHEJ complex showed that XLF bridges two Ku70/80-LIG4/XRCC4 complexes bound on two different DNA ends^20,21^, suggesting a key role of XLF in the end-bridging / synapsis step. How does this step take place in plants? Our results suggest that AtPAXX is likely to play a similar role to XLF by interacting with both AtXRCC4 and AtKu70/80. Since mammalian PAXX and XLF have been shown to have a redundant function, it is unsurprising that one of them is enough for NHEJ. We cannot rule out that plants have other redundant proteins to PAXX. It is intriguing to further investigate how the NHEJ synapsis complex is formed in plants and how this impacts the efficiency of NHEJ.

In summary, our results suggest that NHEJ and its constituent proteins evolved to meet requirements in genome maintenance in different eukaryotes while retaining key conserved functions. This suggests that there is no universal model to explain NHEJ in eukaryotes. Understanding how exactly NHEJ proteins work in different organisms needs detailed experimental studies in these organisms.

## Materials and methods

### Sequence analysis

Homologues of NHEJ proteins were searched using a combination of Psi-Blast^56^ and Jackhmmer. Sequence alignments were generated using ClustalOmega^57^ and Muscle^58^ on Seaview^59^. Phylogenetic trees are created using PhyML^60^ on Seaview and visualised using FigTree. Single chains of AlphaFold models were downloaded from Uniprot, whereas models of homo and heteromultimers were generated using AlphaFold Colab^61,62^.

### Constructs

The open-reading frame of *A. thaliana paxx* was synthesised and codon-optimised for the *E. coli* expression (ThermoFisher Scientific), whereas *A. thaliana lig4* and *xrcc4* genes were gifts from Dr Chris West (University of Leeds). *Atpaxx* and *Atxrcc4* constructs were cloned into pHAT4 and pGAT3 vectors, respectively^63^. *A. thaliana ku70/80* co-expression vector pFastDual-*ku70/80* was a gift from Dr Karel Riha (CEIT, MU)^49^. *A. thaliana lig4* and *xrcc4* genes were cloned into the pFastBac Dual vector.

### Protein purification

AtPAXX fused to a hexahistidine tag (pHAT4-*atpaxx*) was expressed in BL21(DE3) cells (NEB). The proteins were purified using Ni-NTA (Qiagen), and the His tag was cleaved by the Tev protease. Untagged AtPAXX was further purified using a HiTrap Q column (Cytiva). The single peak containing the protein was collected and concentrated using Vivaspin 20 (Cytiva). The protein in a storage buffer (20 mM Tris pH7.4, 150 mM NaCl, 2 mM DTT) was snap-frozen in liquid nitrogen and stored at -80 °C. AtPAXX^1-154^ was also purified in a similar way. AtPAXX^L228E^ fused to hexa-histidine followed by a GST tag (pGAT3-*atpaxx*^*L228E*^) was expressed in BL21(DE3) (NEB) and purified using a 5 ml GSTrap HP column (Cytiva). The His and GST tag were cleaved by the Tev protease and separated from AtPAXX^L228E^ using Ni-NTA (Qiagen). AtPAXX^L228E^ was loaded onto a HiTrap Q HP column equilibrated with 20 mM Tris pH7.4, 50 mM NaCl, 2 mM DTT eluted using a linear gradient of the same buffer but 500 mM NaCl. Peak fractions containing AtPAXX^L228E^ were collected and added to 20 mM Tris-HCl pH7.4, 2 mM to adjust the NaCl concentration to 150 mM based on the chromatography profile of the peak fractions. The protein was concentrated using Vivaspin 20, snap-frozen in liquid nitrogen and stored at -80 °C (Figure S2).

AtXRCC4 fused to a hexa-histidine followed by a GST tag (pGAT3-*atxrcc4*) was expressed in BL21(DE3) cells (NEB), The proteins were purified using GST sepharose (Cytiva), and the his-GST tag was cleaved by the Tev protease. The tag was separated from untagged AtXRCC4 using Ni-NTA. Purified AtXRCC4 was stored in the same final buffer used for AtPAXX and stored at -80 °C after snap-frozen in liquid nitrogen (Figure S2). A shorter version of AtXRCC4^17-163^ was also purified in a similar way (Figure S2).

AtKu70/80 was expressed in Sf9 cells and purified as described previously^34^. The frozen cells were suspended in 50 mM sodium phosphate buffer pH 7.4, 150 mM NaCl, 5% (v/v) glycerol, 2 mM 2-mercaptoethanol, 10 mM imidazole, 20 units of benzonase (Millipore) supplied with cOmplete, EDTA-free protease inhibitor cocktail (Roche) and lysed by sonication. The lysate was incubated for 30 min on ice and diluted 5-fold with 50 mM sodium phosphate buffer pH 8.0, 1M NaCl, 5% (v/v) glycerol, 400 mM ammonium acetate, 2 mM 2-mercaptoethanol, 10 mM imidazole. The cell debris was removed by centrifugation for 15 min at 35,000 g, 4°C. His-tagged AtKu70/80 was isolated from the supernatant using HisPur Ni-NTA resins (Thermo Fisher) and dialysed in 20 mM HEPES pH7.5, 200 mM NaCl, 2 mM 2-mercaptoethanol, 30 mM imidazole overnight after the TEV protease was added to cleave the His tag. Untagged AtKu70/80 was isolated from the His-tag using the Ni-NTA resin and loaded onto a HiTrap Heparin column (Cytiva) equilibrated with the dialysis buffer. AtKu70/80 bound on the column was eluted with 20 mM HEPES pH7.5, 500 mM NaCl, 2 mM DTT, and concentrated and buffer-exchanged to 20 mM HEPES pH7.5, 150 mM NaCl, 2 mM DTT before being snap-frozen in liquid nitrogen to be stored at -80°C (Figure S2).

AtLIG4/XRCC4 (pFastBacDual-*atlig4/xrcc4*) were co-expressed in Sf9 cells, which were suspended in 50 mM Tris-HCl pH7.4, 300 mM NaCl, 2 mM 2-mercaptoethanol, 10 mM imidazole, 5%(v/v) glycerol supplied with cOmplete, EDTA-free protease inhibitor cocktail (Roche). The cells were lysed by sonication, and the cell debris was removed by centrifugation for 30 min at 35,000 g, 4°C. The supernatant was collected and was incubated with HisPur Ni-NTA resins (Thermo Fisher) for 90 min at 4°C. The resins were washed with 50 mM HEPES pH 7.0, 300 mM NaCl, 2 mM 2-mercaptoethanol, 30 mM imidazole, 5%(v/v) glycerol, followed by 50 mM HEPES pH 7.0, 200 mM NaCl, 2 mM 2-mercaptoethanol, 30 mM imidazole, 5%(v/v) glycerol. Bound proteins on the resins were eluted using 50 mM HEPES pH 7.0, 200 mM NaCl, 2 mM 2-mercaptoethanol, 300 mM imidazole, 5%(v/v) glycerol. The eluted proteins were diluted 2-fold using water and loaded onto a 5 ml HiTrap Heparin HP column (Cytiva) equilibrated with 20 mM HEPES pH 7.5, 100 mM NaCl, 2 mM DTT. Bound proteins on the column were eluted using a linear gradient of 20 mM HEPES pH 7.5, 1 M NaCl, 2 mM DTT. Fractions containing AtLIG4/XRCC4 were collected, concentrated and loaded onto Superdex 200 increase 10/300 GL (Cytiva). Fractions containing AtLIG4/XRCC4 were collected and concentrated before being snap-frozen in liquid nitrogen and stored at -80°C (Figure S2).

### X-ray crystallography

Crystals of AtPAXX were obtained using the sitting drop vapour diffusion method where 17 mg/ml of AtPAXX was mixed with 20%(w/v) PEG550MME, 100 mM NaCl, 100 mM Bicine pH9.0 in the 1:1 ration. The crystals were cryo-protected in 33%(w/v) PEG550MME, 100 mM NaCl, 100 mM Bicine pH9.0 and snap-frozen in liquid nitrogen. Their diffraction data were collected at I04 in Diamond Light Source and were processed using their automatic pipeline Dial^64^ and reduced using Aimless^65^ (Table 1). Initial phases of structure factors were obtained by the molecular replacement using the Phaser module in PHENIX suite^66^, using a search model of the AtPAXX dimer generated using AlphaFold Colab^61,62^. The model of AtPAXX was manually and computationally refined using Coot and the phenix.refine module until no further improvement of the electron density map of AtPAXX was observed. TLS groups were selected as each chain, and non-crystallographic symmetry restraints were applied for the model refinement.

### GST-pulldown assay

300 μg of GST-AtPAXX constructs or GST-hPAXX^1-145^ were mixed 20 ul of glutathione sepharose 4B beads (Cytiva) and incubated for 30 min at room temperature in a GST pulldown buffer (20 mM Tris-HCl pH8.0, 200 mM NaCl, 5 mM DTT, 10%(v/v) glycerol, 0.5%(v/v) NP-40. After washing unbound proteins, the beads were suspended in 500 μl of the pulldown buffer and incubated with 100 μg of purified AtXRCC4 for 30 min at room temperature or with lysates of 50 ml cell culture of Sf9 cells expressing AtKu70/80 described above for 60 min at 4°C. After washing beads, TEV protease was added to them to release PAXX constructs and their bound proteins from the beads. Supernatants were then analysed by SDS-PAGE.

### Electrophoretic mobility shift assay

The protocol has been described previously^34^. 20 nM 6FAM-labelled 50bp dsDNA was mixed with 20 nM AtKu70/80 or 200 nM AtPAXX, AtPAXX^1-154^, AtPAXX^L228E^ or different combinations of them as described in the main text in 20 mM Tris-HCl pH7.4, 100 mM KCl, 5% glycerol, 1 mM DTT, 10 μg/ml BSA for 30 min at room temperature. The samples were applied onto a 5% polyacrylamide gel in 0.5x TBE buffer for electrophoresis at 4 °C. DNA was visualised by iBright imager (ThermoFisher Scientific).

### DNA ligation assays

Blunt-ended DNA substrates were prepared by digesting the pGEX6-p1 vector by EcoRV (New England Biolabs) followed by heat inactivation and purification using a gel extraction kit (Qiagen). 50 ng of the DNA were incubated with different combinations of NHEJ proteins (25 nM AtKu70/80, 25 nM AtLIG4/XRCC4, 250 nM AtPAXX, 250 nM AtPAXX^L228E^) in 20 μl of a ligation buffer (25 mM Tris-HCl pH7.4, 100 mM KCl, 1 mM MgCl_2_, 1 mM DTT, 5%(v/v) glycerol, 10 μM ATP, 10 μg/ml BSA). First, the NHEJ proteins except for AtLIG4/XRCC4 were incubated with DNA for 5 min and another 30 min at 37°C after the ligase was added. The reaction was stopped by adding 2 μl of 0.1%(w/v) SDS, 100 mM EDTA and 1 μl of 4 mg/ml Proteinase K (ApexBio Technology) and incubating for 30 min at 50°C. Reaction mixtures were analysed by being applied onto 0.8% agarose gel supplied with SYBR Safe (ThermoFisher Scientific) in TBE buffer for electrophoresis.

## Acknowledgements

T.O. is supported by a startup fund from University of Leeds and BBSRC (BB/V003577/1). We would like to thank Drs Karel Riha for pFastBacDual-Ku70/80 and Chris West for *AtXRCC4* and *AtLIG4* gene, Dr Chi Trinh for his assistance with the diffraction data collection and also Prof. Richard Bayliss, Prof. Brendan Davis, Dr Andrew Blackford and Dr Qian Wu for reading the manuscript.

