## Supplemental data for "Plant PAXX has an XLF-like function and stimulates DNA end-joining by the Ku-DNA ligase IV-XRCC4 complex"

**Table S1**. Accession numbers of NHEJ protein orthologs. Sequences of the listed proteins were used to generate Figure 1. Assession numbers are UniProtKB or, if unavailable, NCBI.

| | KU70 | KU80 | DNA-PKcs | LIG4 | XRCC4 | XLF | PAXX | Artemis | CYREN | APLF | Pol lambda/mu/TdT |
|---|---|---|---|---|---|---|---|---|---|---|---|
| HUMAN | P12956 | P13010 | P78527 | P49917 | Q13426 | Q9H9Q4 | Q9BUH6 | Q96SD1 | Q9BWK5 | Q8IW19 | Q9UGP5; Q9NP87; P04053 |
| MOUSE | P23475 | P27641 | P97313 | Q8BTF7 | Q924T3 | Q3KNJ2 | Q8K0Y7 | Q8K4J0 | Q8BHZ5 | Q9D842 | Q9QXE2; Q9JIW4; P09838 |
| XENLA | Q6AZR6 | Q6DDS9 | Q9DEI1 | Q6GLS1 | A0A1L8I1T7 | A0A1L8ENT6 | A0A8J1LHS2 | A0A1L8GTQ6 | A0A1L8GW76 | A0A1L8G793 | A0A1L8FJD6; A0A8J1MQP2; P42118 |
| CHICK | O93257 | A0A8V0ZIP8 | Q8QGX4 | Q90YB1 | F1NDC0 | F1NVP8 | A0A8V1AIN8 | Q5QJC2 | A0A8V0YFQ4 | E1BU46 | F7B5D1; P36195 |
| DANRE | F1QZ60 | A0A0R4IB13 | E7F4J7 | F1Q9E8 | Q6PBP2 | Q6NV18 | B3DFL2 | Q5RGE5 | A0A0R4IQP0 | A0A8M3AL87 | Q6P0S1; Q7ZUU0; Q5J2Q9 |
| NEMVE | A7RH48 | XP_048590284.1 | A7SPU0 | A7SKL4 | A7RTE3 | A7RY17 | A7RHT8 | A7RK35 | | A7RZU3 | XP_032231870.2 |
| CAEEL | Q9U2D2 | Q21829 | | Q95YE6 | | | O45625 | | | | |
| CRAGI | K1R1F0 | K1Q7H0 | K1Q9H7 | K1R0N9 | K1QHC8 | K1QEJ3 | K1RG06 | K1PT56 | K1RRV0 | K1QVD7 | K1PM26 |
| STRPU | A0A7M7P5E8 | A0A7M7RCH6 | A0A7M7NQ10 | A0A7M7HH42 | A0A7M7PCQ4 | A0A7M7G3W9 | A0A7M7PHI1 | A0A7M7PWX6 | A0A7M7HMK7 | A0A7M7T4X9 | A0A7M7NTT0 |
| AMPQE | A0A1X7U768 | A0A1X7V1U2 | A0A1X7VU01 | A0A1X7VRP5 | A0A1X7VLW7 | A0A1X7TIZ9 | A0A1X7VAM5 | A0A1X7VCC2 | | A0A1X7V5F1 | A0A1X7VDS4 |
| DROME | Q23976 | Q9I7M8 | | Q9VYA5 | Q9VSZ2 | Q9W455 | | | | A8JR14 | |
| BATDJ | F4PDU1 | F4P5J4 | F4PB28 | | | | | | | | |
| YEAST | P32807 | Q04437 | | Q08387 | P53150 | Q06148 | | | | | P25615 |
| SALR5 | F2TX04 | F2UHV6 | F2U190 | F2U9A1 | F2TZY3 | F2UJH2 | F2UPH1 | | | F2UAM4 | F2UQK2 |
| DICDI | Q54MA9 | Q54LY5 | Q54UC0 | Q54CR9 | Q54YJ7 | Q54P54 | | | | | |
| CHLRE | A0A2K3D1K3 | A8ICD8 | A0A2K3DEN6 | A0A2K3DA15 | A0A2K3CU35 | | | | | | A0A2K3CPZ5 |
| ARATH | Q9FQ08 | Q9FQ09 | | Q9LL84 | Q682V0 | | F4KC77 | | | | Q9FNY4 |
| PARTE | Q4LBE8 | A0C319 | A0CIS1 | A0DK55 | A0BK43 | A0CKA0 | A0EGF5 | | | | A0CMJ3 |
| TETTS | A4VEG9 | I7LWU7 | Q22NE4 | Q23RI5 | W7XIC1 | I7MIB7 | I7MJ21 | | | | |
| NAEGR | D2W5B9 | D2VUC7 | D2V5J3 | D2VDE2 | D2VM57 | D2VLF4 | D2VAC1 | | | | |
| GIAIA | | | | | | | | | | | |
| TRYCC | Q4CY05 | Q4DHP5 | Q4DW48 | | | | Q4E664; Q4E006 | | | | |
| LEIMA | E9ADW3 | Q4Q7S5 | Q4Q1C5 | | | | Q6S998 | | | | |
| PHYPR | W2KE29 | W2KQ35 | W2LI30 | W2LDA3 | W2YUL7 | | | | | | W2LF93 |

**Table S2.** PAXX orthologs in streptophyta. Assession numbers are Assession numbers are UniProtKB or, if unavailable, NCBI.

| **Organism** | **Assession number** |
|---|---|
| *Oryza sativa Japonica* | A0A8J8XRR6 |
| *Aristolochia fimbriata* | KAG9451149.1 |
| *Amborella trichopoda* | U5D7E4 |
| *Ceratopteris richardii* | A0A8T2UUR8 |
| *Physcomitrium patens* | A0A7I4EPE9 |
| *Selaginella moellendorffii* | D8RQ89 |
| *Klebsormidium nitens* | A0A0U9HJC7 |

### Figure S1

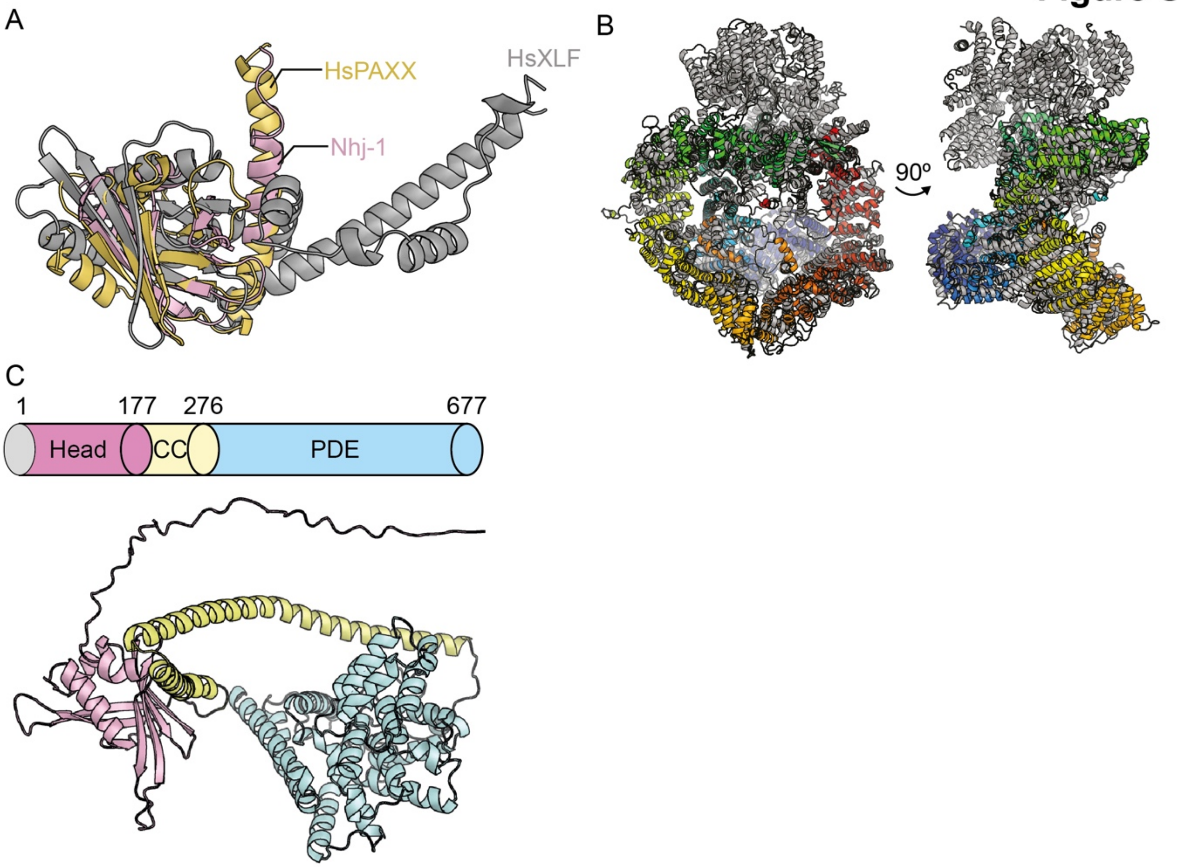


**Figure S1**. AlphaFold models of NHEJ orthologs. (A) Nhj-1 is PAXX ortholog. An AlphaFold model of Nhj-1 (pink) is compared with crystal structures of human PAXX and XLF (PDB codes: 3WTD (yellow) and 2QM4 (grey)) by superimposing the head domains of the structures. (B) *Trypanosome* putative DNA-PKcs lacks the kinase domain. An AlphaFold model of a putative DNA-PKcs ortholog is compared with the crystal structure of human DNA-PKcs (PDB code: 5LUQ (grey)). Trypanosome DNA-PKcs is presented in a gradient of blue (N-terminus) and red (C-terminus). (C) Trypanosome XRCC4/XLF/PAXX has a PDE domain. A schematic domain diagram shows the domain arrangement of a putative ortholog of XRCC4/XLF/PAXX in *Trypanosoma*. An AlphaFold model of the protein is shown below the diagram with the same domain colour scheme.

**Figure S2**

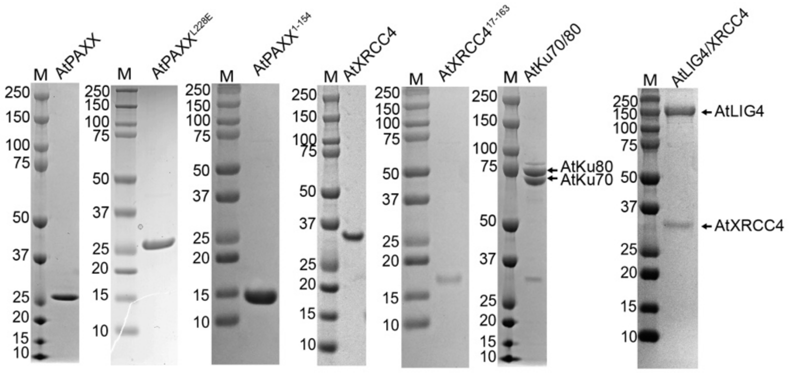


**Figure S2**. Purified recombinant Arabidopsis NHEJ proteins. 2 μg of each purified protein were loaded onto an SDS-PAGE gel. "M" indicates protein ladders, of which molecular weights (kDa) are shown on the left.
